# CFD-based Bayesian Optimization of Stirring Strategies in Stirred Tank Cultures of Pluripotent Stem Cell Spheroids

**DOI:** 10.64898/2026.07.06.735037

**Authors:** Ikki Horiguchi, Kippei Okada, Yasunori Okano

## Abstract

The suspension culture of pluripotent stem (PS) cells in stirred bioreactors poses a delicate balance between maintaining homogeneous cell dispersion and avoiding excessive shear stress that can compromise cell viability and pluripotency. In this study, we used computational fluid dynamics (CFD) coupled with a discrete particle method (DPM) to simulate iPS cell behavior in a 5 mL delta-impeller stirred tank. Our analysis revealed that upward flow at the tank bottom and downward flow at the top are critical for maintaining a stable suspension. To optimize the stirring protocol, we applied Bayesian optimization to identify a time-dependent stirring schedule that begins with a high-speed phase for resuspension, followed by a low-speed phase for sustained suspension with minimal hydrodynamic stress. The optimized schedule demonstrated improved floating rate and reduced slip velocity, indicating lower mechanical stress on cells. These findings provide engineering insights into scalable bioreactor operation, contributing to the design of robust iPS cell manufacturing systems.

**Highlights:**

- CFD-DPM simulations predicted iPS spheroid motion in a stirred tank.
- Bayesian optimization identified a two-step agitation strategy.
- High initial agitation promoted rapid particle resuspension.
- Low subsequent agitation maintained suspension with lower downflow.
- Particle Reynolds number correlated with the floating rate.

## 1. Introduction

Induced pluripotent stem (iPS) cells are promising cell sources for regenerative medicine, drug discovery, and disease modeling due to their ability to differentiate into various cell types and proliferate indefinitely [1]. As clinical and industrial applications advance, the need for scalable, efficient, and controllable suspension culture systems for iPS cells is rapidly increasing. For example, stirred tank bioreactors have become the predominant platform for large-scale expansion of iPS cells due to their scalability, homogeneity, and compatibility with automated systems [2, 3].

However, suspension cultures of iPS cells present significant engineering challenges. Insufficient agitation causes cell sedimentation and non-uniform spheroid formation and growth, while excessive agitation generates hydrodynamic shear stress, potentially damaging cell membranes or altering differentiation capacity [4, 5]. Therefore, an optimized stirring strategy is required to ensure sufficient suspension culture without imposing harmful mechanical stress on the cells. However, iPS cell culture requires high cost and a long culture period, which limits the number of trials for experimental optimization.

Therefore, various analysis methodologies have been attempted based on experimental and computational strategies. In an experimental strategy, visualization of medium flow and cell trajectories [6, 7] was developed for the analysis of suspension culture process, but there is a limitation in visualization of cell behavior due to the size difference between the bioreactor and cells. In terms of computational strategy, computational fluid dynamics (CFD) has been widely used to simulate hydrodynamic environments in bioreactors, providing information on local shear distributions and flow patterns[8, 9]. CFD studies of bioreactors have frequently been limited to the analysis of hydrodynamic behavior within the reactor. However, in the design of suspension culture processes for animal cells, such as human induced pluripotent stem cells (hiPSCs), it is crucial to understand how the flow environment influences the cells themselves. In particular, flow-induced factors such as shear stress can significantly affect cell growth, viability, and differentiation, and should therefore be considered in bioreactor design. The discrete particle method (DPM), when coupled with CFD, enables the modeling of particle–fluid interactions at the microscale [10], allowing simulation of the behavior of cell aggregates or microcarriers in suspension [11]. Despite their utility, these simulations are computationally demanding, making manual tuning of stirring schedules inefficient.

To address this, we propose a framework that combines CFD–DPM simulation with Bayesian optimization to identify optimal time-dependent stirring schedules. Bayesian optimization is widely used for optimizing expensive black-box functions [12] with a limited number of evaluations including CFD [13, 14]. This approach allows for efficient exploration of parameter space with limited computational cost [15], guiding the design of stirring profiles that improve suspension while minimizing shear-related damage. In this study, we apply this framework to a small-scale stirred tank system and evaluate its effectiveness in maintaining particle suspension under realistic culture conditions. Through the evaluation, we also suggest the key phenomena to keep cells floating with low shear stress.

## 2. Numerical methodologies

### 2.1 Model overview

The commercially available disposable small-scale bioreactor with delta-shaped paddle was applied as a stirred tank reactor model for calculation (Fig. 1). Agitation rate (*ω*) was the parameter for optimization. The working volume was set to 5 mL to reduce computational cost. OpenFOAM-v2106 was used for developing and running solvers to calculate the flow dynamics in the stirred-tank bioreactor. Calculation was performed by the Intel(R) Xeon(R) Gold 6338 CPU @ 2.00GHz. The computation time was approximately 100 h.

**Figure 1.**
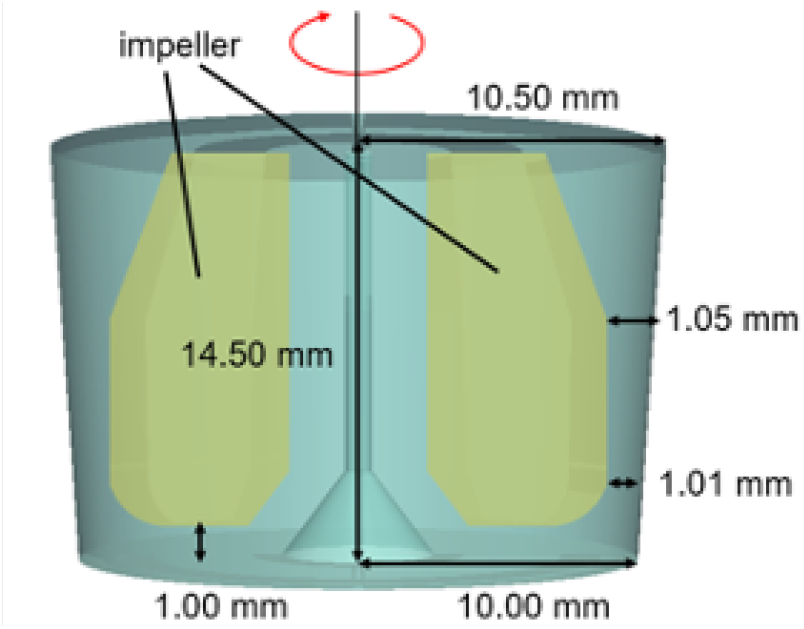
Geometrical model of a small-scale stirred tank bioreactor for CFD analysis.

### 2.2 Governing equations

#### 2.2.1 Fluid phase

Culture medium was assumed to be a Newtonian fluid and incompressible. Flow velocity and pressure distribution was determined by solving the following continuity equation and the Navier-Stokes equation. The medium flow in the reactor was assumed to be laminar flow.

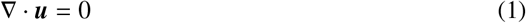

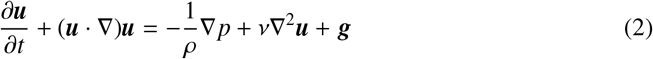

#### 2.2.2 Solid phase

The cells in the bioreactor were considered solid spheres and the numerical methodology was based on previous reports. The particle density (*ρ*_p_) was 1.075×10^3^ kg/m^3^ and the diameter (*d*_p_) was 1.0×10^−4^ m, which is assumed as a diameter of iPS cell aggregates in early stage of culture. The movement of the cells was calculated by discrete particle method (DPM). The spheres are assumed to obey Newton’s equations of motion.

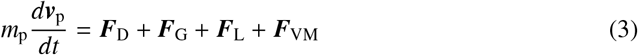

The right-hand side represents, respectively, the drag force from the fluid, the gravity considering buoyancy, the lift force due to the pressure difference around the particle, and the virtual mass force.

In this model, particle–particle collisions were ignored because the particle concentration was sufficiently dilute, resulting in a low probability of interparticle interactions.

### 2.3 Bayesian optimization

For Bayesian optimization of the agitation parameters; initial agitation rate (*ω*_1_), secondary agitation rate (*ω*_2_) and switching time (*t*_s_) of stirred tank bioreactors. To consider actual bioreactor operations, a switching period of 2 seconds was implemented for switching agitation speed. Accordingly, the switching time was defined as a point at which the switching agitation speed is fully completed. The objective function was defined below as previously reported.

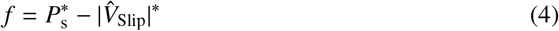

Here, *P*^*^_s_ represents the floating rate, defined as the numerical ratio of cell particles excluding those located in the meshes adjacent to the bottom surface, which are considered to be settled. 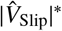 represents the ensemble average of the relative velocity of the particles with respect to the fluid. The starred parameters are normalized to the maximum value. The input parameter sets to be optimized are listed in Table 1.

**Table 1.**
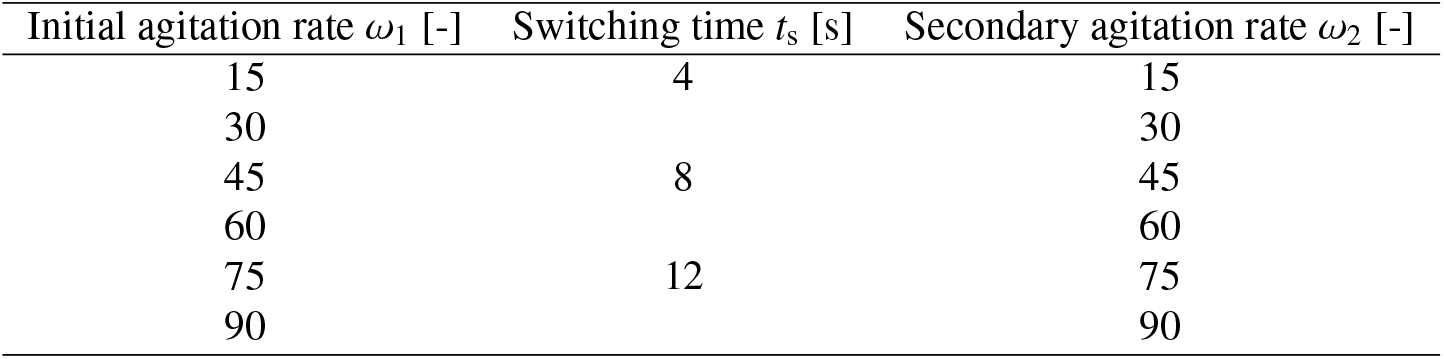
Input agitation parameters to be optimized Bayesian optimization.

| Initial agitation rate $\omega_1$ [-] | Switching time $t_s$ [s] | Secondary agitation rate $\omega_2$ [-] |
| --- | --- | --- |
| 15 | 4 | 15 |
| 30 |  | 30 |
| 45 | 8 | 45 |
| 60 |  | 60 |
| 75 | 12 | 75 |
| 90 |  | 90 |

### 2.4 Dimensionless analysis

For further understanding and generalizing the phenomena governing the interactions between the particles and the fluid, the Navier-Stokes equation was nondimensionalized as Eq. (5).

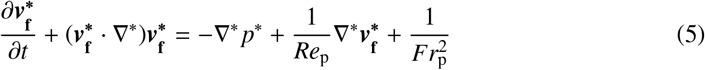

Here, particle Reynolds number *Re*_p_ and particle Froude number *Fr*_p_ were defined as

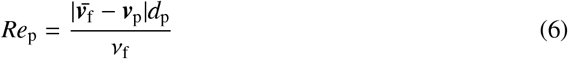

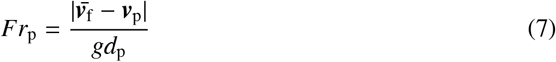

## 3. Results and discussion

### 3.1. Analysis of medium flow and cell behavior in stirred tank

To understand the behavior of cell particles within the reactor, we first examined the effect of agitation speed on the floating rate after a fixed period of operation. For this evaluation, we utilized two distinct initial cell distributions: one where cells were localized at the bottom and another where they were suspended at the top. Fig. 2 illustrates the changes in two distinct suspension rates across different stirring speeds: the rate for particles initially in a floating state and the final suspension rate for particles initially settled at the bottom. These results suggest that while a high stirring speed is effective for resuspending particles from the bottom, it concurrently promotes the sedimentation of particles that are already in suspension.

**Figure 2.**
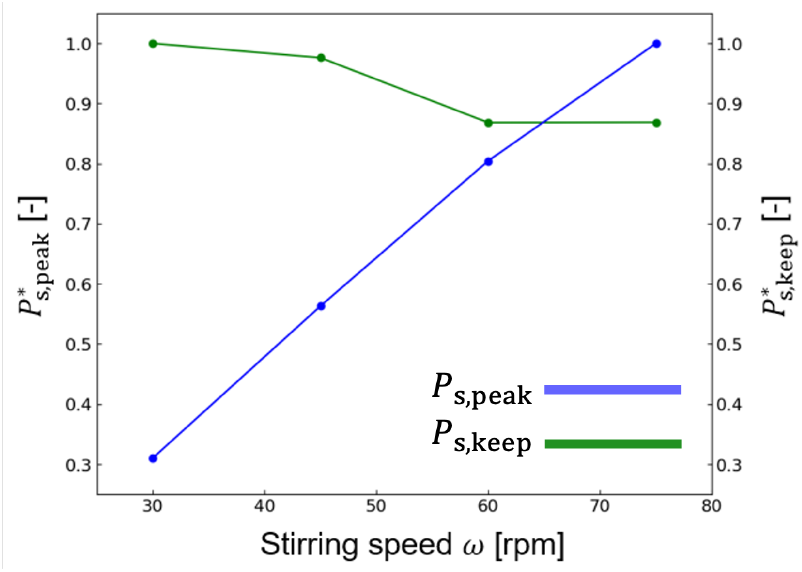
Relationship between the normalized floating rate (scaled to a maximum of 1) and agitation speed: (Green) normalized floating rate of cells initially in a suspended state *P*_*s,keep*_; (Blue) floating rate after 30 seconds for cells initially placed at the bottom *P*_*s,peak*_.

Fig. 3 visualizes (A) the direction of liquid flow and (B) the time-average intensity of the upward flow inside the reactor. It can be observed that at a low agitation speed, such as 30 rpm, a circulatory flow develops in the vertical direction; conversely, at a high agitation speed, such as 75 rpm, a strong liquid flow toward the wall surface is generated. Considering the motion of particles carried by the flow, low agitation speeds likely maintain suspension through vertical vortices. Conversely, high agitation speeds suggest a tendency for particles to be driven toward the wall and settle along its surface. Analysis of the time-averaged upward flow shows that high agitation speeds promote a strong downward flow in the upper reactor, likely driving cells toward the bottom. Conversely, at lower speeds, the absence of this downward flow helps maintain cell suspension.

**Figure 3.**
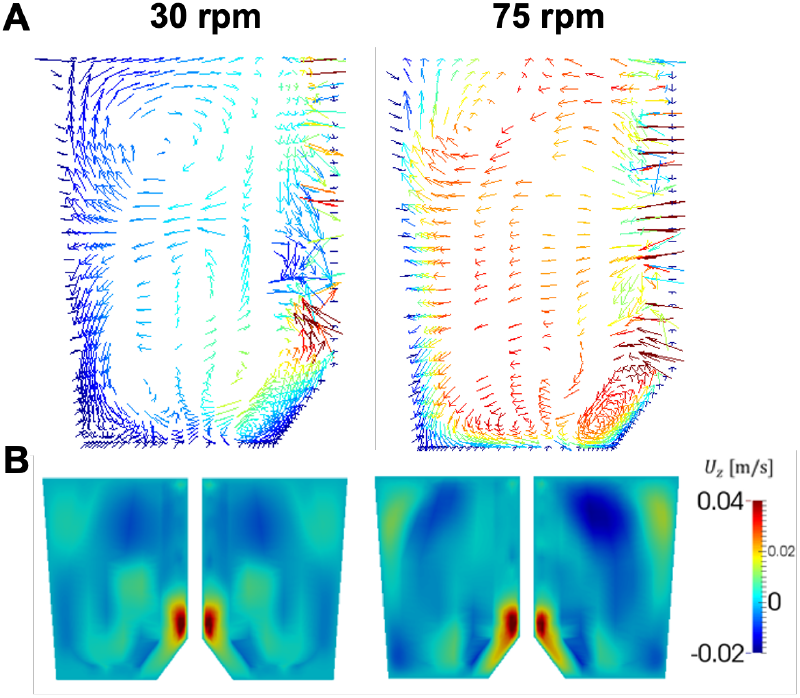
Flow velocity distribution within the stirred bioreactor at high (75 rpm) and low (30 rpm) stirring speed: (A) two-dimensional flow vector distribution; (B) upward flow distribution.

The results above indicate that while high agitation speeds are advantageous for lifting particles from the bottom, they also promote the settling of suspended particles due to strong downward and wall-directed flows. Conversely, low agitation speeds are disadvantageous for particle lift-off but are better suited for maintaining the suspended state. These findings suggest that precise control of agitation speed is crucial for maintaining cells in suspension.

### 3.2 Bayesian optimization of agitation schedule

According to the repeated numerical simulations conducted by Bayesian optimization (Fig. 4), the highest value of the objective function was achieved with the parameters, (*ω*_1_, *t*_*s*_, *ω*_2_)=(60, 4, 15), suggested during the fifth iteration. The proposed protocol involves starting with a high agitation speed and subsequently shifting to a lower speed; this approach is consistent with the observations from the CFD analysis regarding particle behavior and flow patterns.

**Figure 4.**
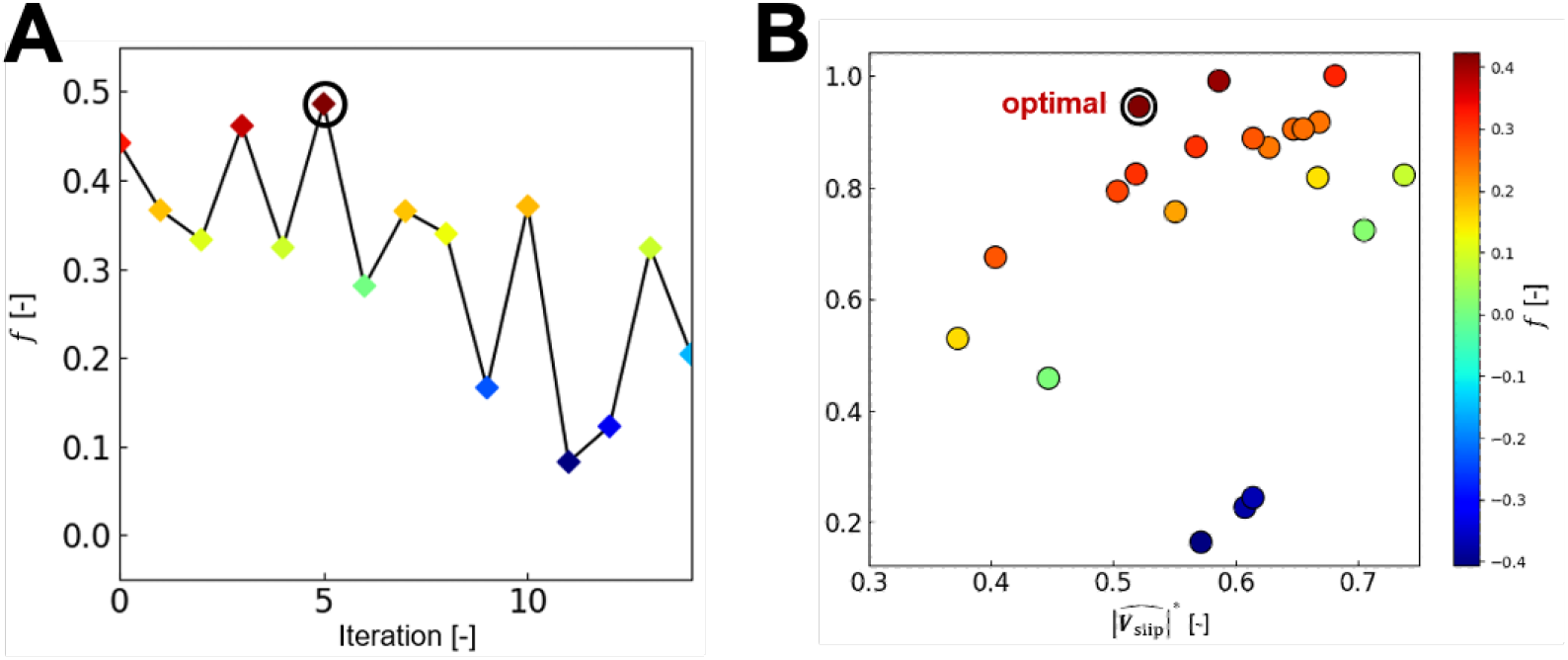
Result overview of Bayesian optimization: (A)profiles of objective function during iterations; (B)mapping of conditions and objective functions.

Since the optimal parameters include minimum value in switching time and secondary agitation rate thus the ideal operating condition may be below search range. However, considering practical factor such as the agitation torque and the time required to reach setpoints in an actual bioreactors, it is deemed preferable to adopt the optimal values identified within the current search range.

These practical factors must also be considered when applying optimization to actual bioreactor operations. For example, while the agitation rate in numerical simulations can be switched instantaneously, in practice, it is adjusted gradually. Such challenges in bioreactor optimization require an integrated approach considering not only fluid dynamics but also mechanical limitations.

### 3.3 Evaluation of optimized agitation schedule

Fig. 5 illustrates time profiles of floating rate(A) and relative velocity(B) in constant agitation conditions(75 and 30 rpm) and optimal condition,(*ω*_1_, *t*_*s*_, *ω*_2_)=(60, 4, 15). In high-speed condition (75 rpm), floating rate increased rapidly in the early stage but gradually declined after reaching its peak. Conversely, in low-speed condition (30 rpm), while the initial increase was limited, the floating rate remained stable at a later stage. In contrast, the optimal condition achieved a rapid initial increase in floating rate and successfully maintained a high value, nearly at its peak level, throughout the operation.

**Figure 5.**
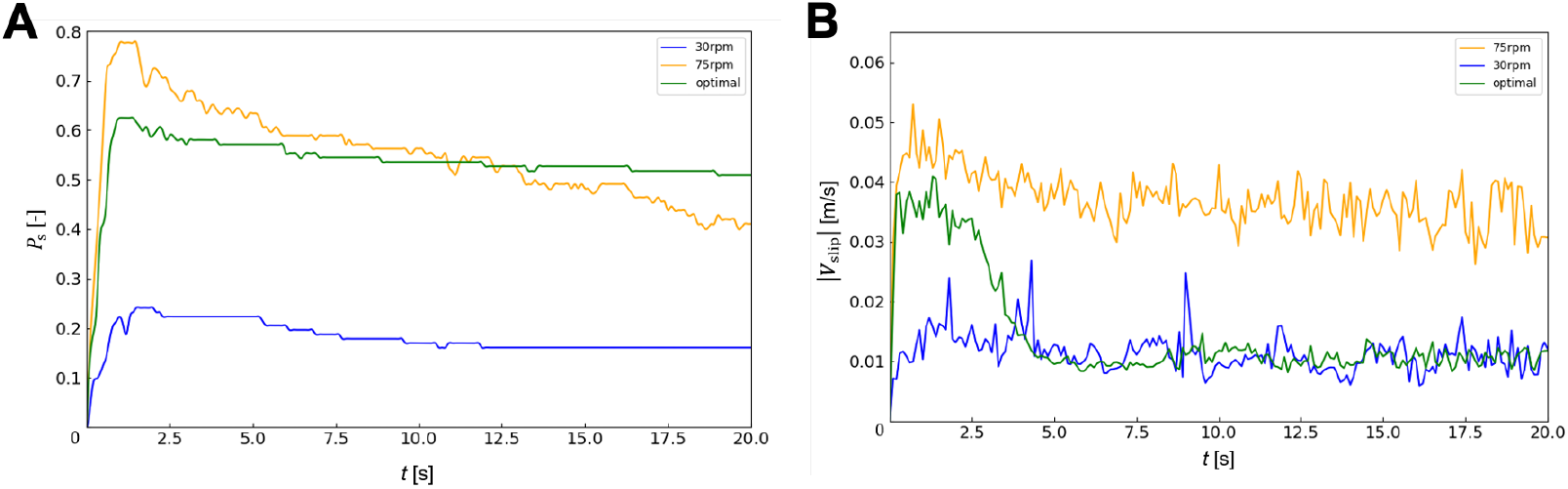
Time profiles of (A)floating rate and (B) relative velocity between medium and particles in static condition (75 and 30 rpm) and optimized condition,(*ω*_1_, *t*_*s*_, *ω*_2_)=(60, 4, 15).

Relative velocity remained stable under constant agitation conditions(Fig. 5B). In contrast, the optimal condition exhibited high relative velocity at the initial period and the gradual decrease during the agitation speed transition (from 2 to 4 seconds) and stable low level as 30 rpm.

These results demonstrate that the optimized operating conditions successfully apply high shear stress during the initial stage, which is necessary for cell lift-off, while minimizing shear stress during the subsequent long-term operation. This finding highlights the importance of time-dependent control of agitation speed for achieving both efficient resuspension and stable suspension, providing a useful strategy for the design of bioreactor operation for shear-sensitive cells.

However, several limitations of this study should be noted. First, due to the high computational cost of CFD–DPM simulations, the present analysis was restricted to a small-scale system (5 mL). As the number of computational cells and particles increases significantly with reactor size, simulation of larger-scale systems becomes computationally prohibitive. Therefore, no direct validation at larger scales was performed, and the extrapolation of the present results to industrial-scale bioreactors remains an important challenge. In particular, changes in key dimensionless parameters, such as the Reynolds and Froude numbers, may alter the flow structure and particle dynamics, potentially affecting the applicability of the proposed strategy.

In addition, experimental validation in actual cell culture systems is limited. Direct observation of sedimentation behavior and particle trajectories in three-dimensional and opaque culture environments is technically challenging, making it difficult to establish a direct causal relationship between simulated hydrodynamic parameters and observed culture outcomes. Future work should therefore focus on linking simulation-derived indicators, such as floating rate and slip velocity, with experimentally measurable parameters including cell viability, aggregation behavior, and differentiation potential.

This limitation highlights the need for a generalized framework that enables comparison and interpretation of particle suspension behavior independent of system scale and operating conditions.

### 3.4 Effect of dimensionless numbers on floating rate

Particle suspension in stirred vessels has traditionally been discussed in terms of the just-suspended speed and dimensionless correlations involving particle properties, impeller geometry, and hydrodynamic conditions [16]. Subsequent studies have further investigated the mechanisms of complete suspension and the effects of operating and geometrical parameters on the minimum agitation speed required for particle suspension [17, 18]. However, the applicability of empirical just-suspension correlations can be limited when the vessel geometry, impeller configuration, particle properties, or operating regime differ from the conditions used to derive the correlations [19]. Therefore, in the present study, we performed a dimensionless analysis using the particle Froude number (*Fr*_*p*_) and particle Reynolds number (*Re*_*p*_) to identify the key parameters governing the suspension behavior of cell particles in the stirred tank bioreactor.

According to the scatter plot between floating rate and particle Froude number(Fig. 6A), only a weak correlation was observed between floating rate and particle Froude number. This result suggest that the effect of gravity on the floating rate was limited and drag force generated by the fluid flow predominantly influenced the behavior of the cell particles in the bioreactor.

**Figure 6.**
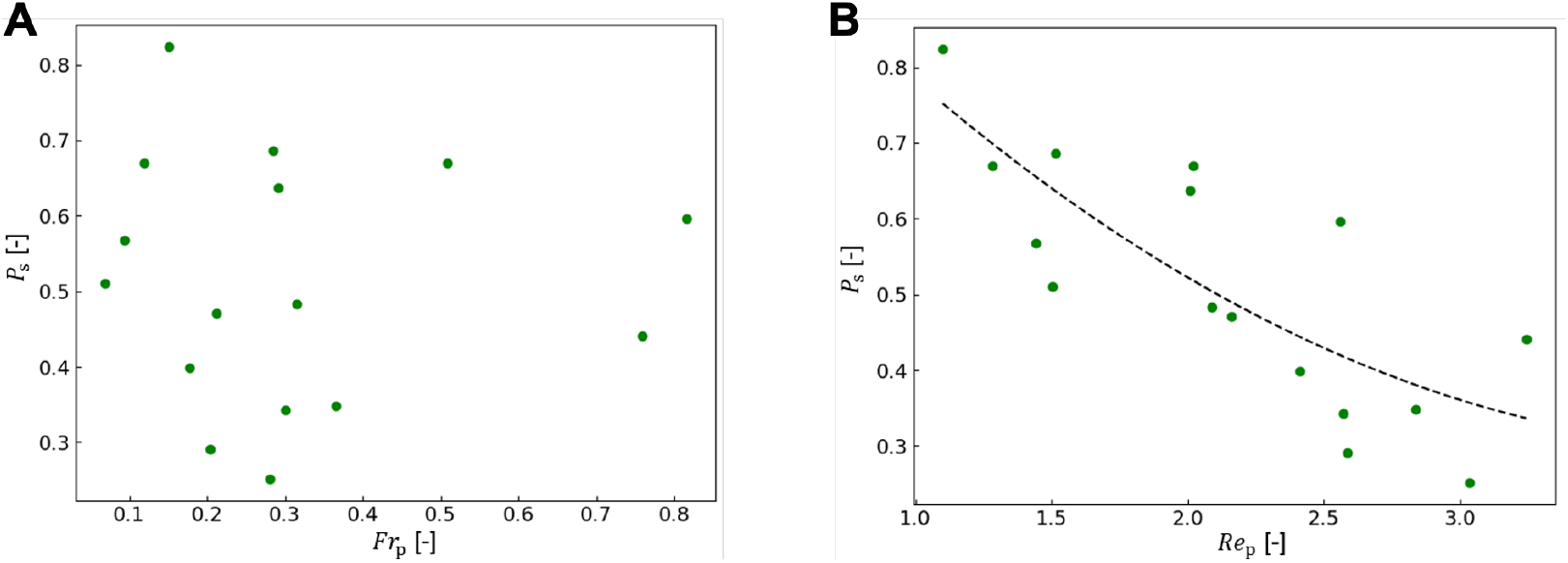
Relation between floating rate and dimensionless numbers around particles; (A) particle Froude number and

In contrast, a negative correlation was observed between the particle Reynolds number and the floating rate. This result suggests that the sedimentation behavior of the cell particles was affected by the balance between viscous and inertial force of fluid field surrounding cell particles. Previous literature suggests that the Reynolds number influences the lift force[20], which is considered to support the findings of this study that the particle Reynolds number impacts the suspension rate.

The results indicate that the floating rate decreased with increasing particle Reynolds number. Although the calculated particle Reynolds numbers under all conditions suggested that the flow around the particles remained in the laminar regime, a higher contribution of inertial forces may reduce the ability of the particles to follow the fluid flow, thereby promoting sedimentation due to inertial effects.

Because the particle Reynolds number is affected by particle size, the growth of cell aggregates accompanying cell proliferation is expected to increase the aggregate diameter. Therefore, as the cells grow, operational conditions may need to be controlled to maintain an appropriate particle Reynolds number. Specifically, this could include reducing the agitation speed and controlling the viscosity of the culture medium.

This result suggests that maintaining an appropriate particle Reynolds number could serve as a useful guideline for adjusting operating conditions during scale-up, even when absolute flow conditions differ.

## Conclusion

In this research, we evaluated CFD-DPM analysis in stirred tank reactor for iPS cell culture and Bayesian optimization was performed to optimize switching agitation rate for further stable cell culture. High stirring speeds were beneficial during the initial phase to resuspend cell particles at the bottom of the bioreactor. However, the analysis suggests that stable suspension after initial phase can be maintained by a lower stirring speed. This strategy enables the maintenance of sufficient upward flow to prevent sedimentation while minimizing the shear stress and the strong downward flow generated by vigorous agitation, which may otherwise promote cell damage and particle settling. This finding provides crucial insights not only for iPS cells but also for the process design of agitated culture systems for human and animal cells that are sensitive to shear stress and require maintenance in a suspended, mixed state.

## Acknowledgement

This research was supported by JSPS KAKENHI Grant number JP23K04507 and JP26K01311.

